# Astrocyte Transcriptomics in a Three-Dimensional Tissue-Engineered Rostral Migratory Stream

**DOI:** 10.1101/2025.02.24.639941

**Authors:** Michael R. Grovola, Erin M. Purvis, Andrés D. Garcia-Epelboim, Elizabeth N. Krizman, John C. O’Donnell, D. Kacy Cullen

**Affiliations:** Center for Neurotrauma, Neurodegeneration & Restoration, Corporal Michael J. Crescenz VA Medical Center, Philadelphia, PA; Center for Brain Injury & Repair, Department of Neurosurgery, Perelman School of Medicine, University of Pennsylvania, Philadelphia, PA; Department of Neuroscience, Perelman School of Medicine, University of Pennsylvania, Philadelphia, PA; Department of Physics and Astronomy, School of Arts and Sciences, University of Pennsylvania, Philadelphia, PA; Department of Bioengineering, School of Engineering and Applied Science, University of Pennsylvania, Philadelphia, PA

**Keywords:** astrocytes, transcriptomics, rostral migratory stream, tissue engineering

## Abstract

The glial tube is a longitudinal structure predominantly composed of densely bundled, aligned astrocytes that projects from subventricular zone (SVZ) to olfactory bulb. Neural precursor cells (NPCs) generated in the SVZ migrate through this glial tube – referred to as the rostral migratory stream (RMS) – to replace olfactory bulb interneurons in the mammalian brain. RMS astrocytes have distinct morphological and functional characteristics facilitating their unique purpose as an endogenous living scaffold directing NPC migration and maturation. However, the transcriptomic factors underlying these unique structure-function attributes versus standard stellate astrocytes have not been examined. We previously developed biofabrication techniques to create the first tissue-engineered rostral migratory stream (TE-RMS) that replicates key features of the glial tube *in vivo*. We have shown that TE-RMS astrocytes exhibit elongated nuclei, longitudinally aligned intermediate filaments, and enrichment of key functional proteins – cytoarchitectural and surface features characteristic of native RMS astrocytes. In the current study, we performed RNAseq on TE-RMS astrocytes in comparison to planar astrocyte cultures to identify gene expression patterns that may underlie their profound morphological and functional differences. Remarkably, we found 4008 differentially expressed genes in TE-RMS astrocytes, with 2076 downregulated (e.g. LOC690251, *ccn5*) and 1932 upregulated (e.g. *lrrc45*, *cntn1*) compared to planar astrocytes. Moreover, there were 256 downregulated and 91 upregulated genes with >3-fold change. We also conducted analyses of gene sets related to cytoskeleton and nuclear structure, revealing greatest enrichment of actin-related components. Overall, the TE-RMS offers a platform to study interplay between transcriptomic and cytoarchitectural dynamics in a unique astrocyte population.

## Introduction

Astrocytes are the most abundant cell type in the mammalian brain and are critical to maintaining homeostasis due to myriad roles including providing metabolic, trophic, and physical support for neurons, controlling the blood-brain barrier, and regulating the synaptic microenvironment (1–4). Astrocytes derive their name from their stereotypical star-shaped morphology and are generally distributed homogeneously in non-overlapping domains throughout the brain (1,5,6). In contrast, the glial tube is composed of a curious astrocyte subtype marked by aligned, bidirectional, interwoven processes in a dense, rope-like bundle (7–10).The glial tube projects longitudinally from the subventricular zone (SVZ) to the olfactory bulb and plays a key role in guiding the migration, maturation, and differentiation of new neurons (11–14).

In particular, the formation of new neurons in the brain, also known as neurogenesis, occurs in the SVZ and continues throughout adulthood in most mammals (15,16). In the SVZ, neural precursor cells (NPCs) are generated and can differentiate into neuroblasts that migrate long distances along the glial tube – referred to as the rostral migratory stream (RMS) – to the olfactory bulb, where they integrate into the existing olfactory circuitry as various types of interneurons (13,14,17,18). Neuroblasts can be diverted from the RMS pathway towards injured brain regions guided by chemo-attractive factors, and can mature into functional neurons of a region-specific phenotype (19–24). Furthermore, functional recovery was found after experimentally enhancing the delivery of neuroblasts from the SVZ into injured brain regions (23–33). Our research team and others have augmented this redirection of neuroblasts utilizing a variety of tissue engineering and biomaterial strategies (34–36).

Specifically, our team developed the first tissue engineered RMS (TE-RMS), a living, biologically-active scaffold comprising aligned, bundled astrocytes that mimics the endogenous RMS. The TE-RMS has been designed to facilitate migration of NPCs to target brain regions (9–12,37,38). Our previous experiments have demonstrated that the TE-RMS can be fabricated from rat astrocytes or human astrocytes derived from adult gingiva, that TE-RMS astrocytes express key functional proteins of the endogenous rat RMS (e.g., ezrin and robo2), and that they facilitate migration of immature rat cortical neurons *in vitro* as well as *in vivo* post-implantation (12). Recently, we have also demonstrated that the TE-RMS can direct migration of neuroblasts specifically harvested from the rat subventricular zone (38). By structurally and functionally mimicking the endogenous RMS, TE-RMS implants may create a migration pathway to injured brain tissue and thus provide a mechanism for gradual yet sustained neuronal replacement as a regenerative medicine strategy.

TE-RMSs are fabricated within hydrogels using geometric cues (e.g., hollow microcolumns or microchannels) and an extracellular matrix coating (Figure 1A-B) (9–12,36,38). This fabrication process induces the self-assembly of dissociated (spherical) astrocytes into three-dimensional longitudinally aligned bundles (Figure 1C). We have previously shown that the TE-RMS replicates the structure and key protein expression of the endogenous rat RMS. We have also quantified the extent of morphological changes in the bundled, longitudinally-aligned TE-RMS astrocytes versus astrocytes in typical planar culture. Specifically, cytoskeletal rearrangement and nuclear elongation were clearly evident in TE-RMS astrocytes, similar to RMS astrocytes *in vivo* (37). Relatively recent scientific advances have identified molecular mechanisms initiated by mechanical stress on the cytoskeleton, which may alter nuclear structure and chromosome dynamics (39,40). These cytoarchitectural adaptations as cells move from two-dimensional environments into three-dimensional environments are necessary for the cell body and nucleus to reshape, allowing cell migration through the available space (41).

**Figure 1.**
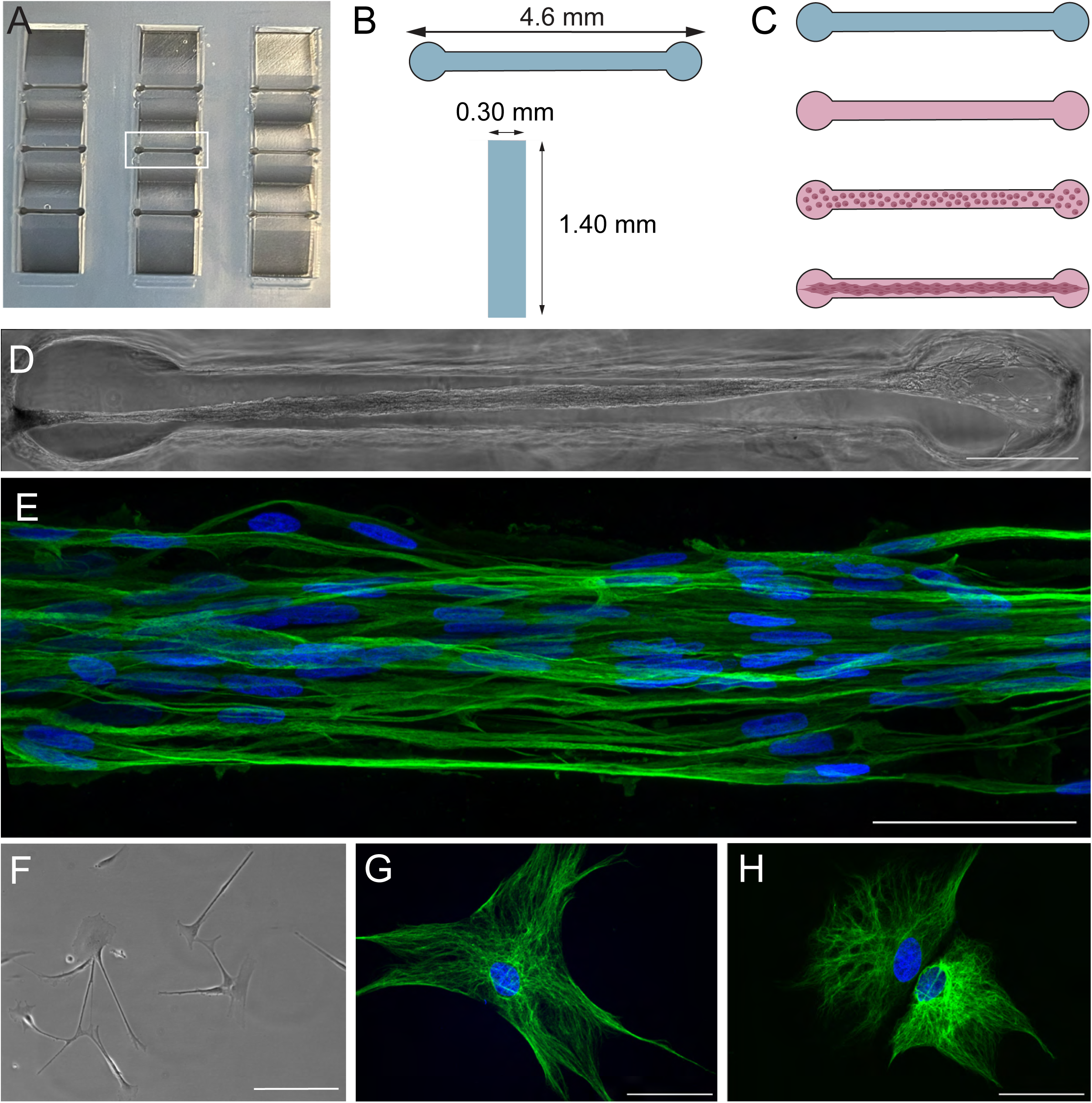
Fabrication and architecture of tissue-engineered rostral migratory streams. TE-RMSs were fabricated in 3×3 grids of hydrogel rectangular microchannels (A). The central microchannel in the 3×3 grid is squared in white (A). Microchannels were 4.6 mm long, 0.3 mm wide, and 1.4 mm deep (B). To fabricate TE-RMSs, microchannels were dried and coated with extracellular matrix. Following complete polymerization and drying of extracellular matrix, microchannels were seeded with a dense suspension of astrocytes that bundled into TE-RMSs (C). Example of a bundled TE-RMS in a microchannel seen under phase microscopy (D). High magnification confocal imaging reveals the structure of TE-RMS astrocytes with elongated cell nuclei and bidirectional intermediate filament processes (E). TE-RMS astrocytes (D-E) have distinct architecture compared to planar astrocytes (F-H). Planar astrocyte samples visible under phase (F) and fluorescence microscopy (G-H) depict astrocytes with round nuclei and intermediate filament processes spread in all directions. Scale bars: 500 microns (D), 20 microns (F), 50 microns (E, G, H).

It is evident that RMS astrocytes have distinct morphological and functional characteristics facilitating their unique purpose as an endogenous living scaffold directing NPC migration and maturation. However, the transcriptomic factors underlying these unique structure-function attributes versus standard stellar astrocytes have not been examined. Accordingly, in the current study we conducted high-throughput RNA sequencing on TE-RMS microtissues to identify gene expression changes that may have contributed to or resulted from morphological changes and compared expression levels to planar astrocytes *in vitro*. Then we performed gene set analysis on annotated gene sets related to cytoskeleton and nuclear structure. We hypothesized that TE-RMS astrocytes would have significant changes to cytoskeletal and nuclear related genes compared to planar astrocytes due to their altered cytoarchitecture. This study provides insight into the transcriptomic and structural responses of astrocytes following TE-RMS formation.

Moreover, this study highlights the TE-RMS as a novel platform to investigate cytoskeletal arrangement and nuclear morphology while also identifying genetic targets to customize astrocyte populations for potential utility in regenerative applications.

## Methods

### Isolation and culture of astrocytes

All described procedures adhere to the National Institutes of Health Guide for the Care and Use of Laboratory Animals and were approved by the Institutional Animal Care and Use Committee at the University of Pennsylvania. Primary cortical astrocytes were harvested from postnatal day 0-1 Sprague-Dawley rat pups (female or male) (Figure 2A). Following dissociation [described previously in (10)], astrocytes were cultured in Dulbecco’s Modified Eagle Medium F12 (DMEM/F12; Gibco #11330032) supplemented with 10% Fetal Bovine Serum (Sigma #F0926) and 1% Penicillin-Streptomycin (Gibco #15140122) antibiotics in a cell culture incubator maintained at 37° C and 5% CO_2_. Astrocyte flasks were passaged with trypsin-EDTA (Gibco #25200056) at 80% confluency to maintain astrocyte cell lines. Passage 3 astrocytes were split into 3 separate passage 4 sister flasks (Figure 2B-C). Astrocytes at passage 4 were used for TE-RMS fabrication and planar astrocyte plating (Figure 2C-E).

**Figure 2.**
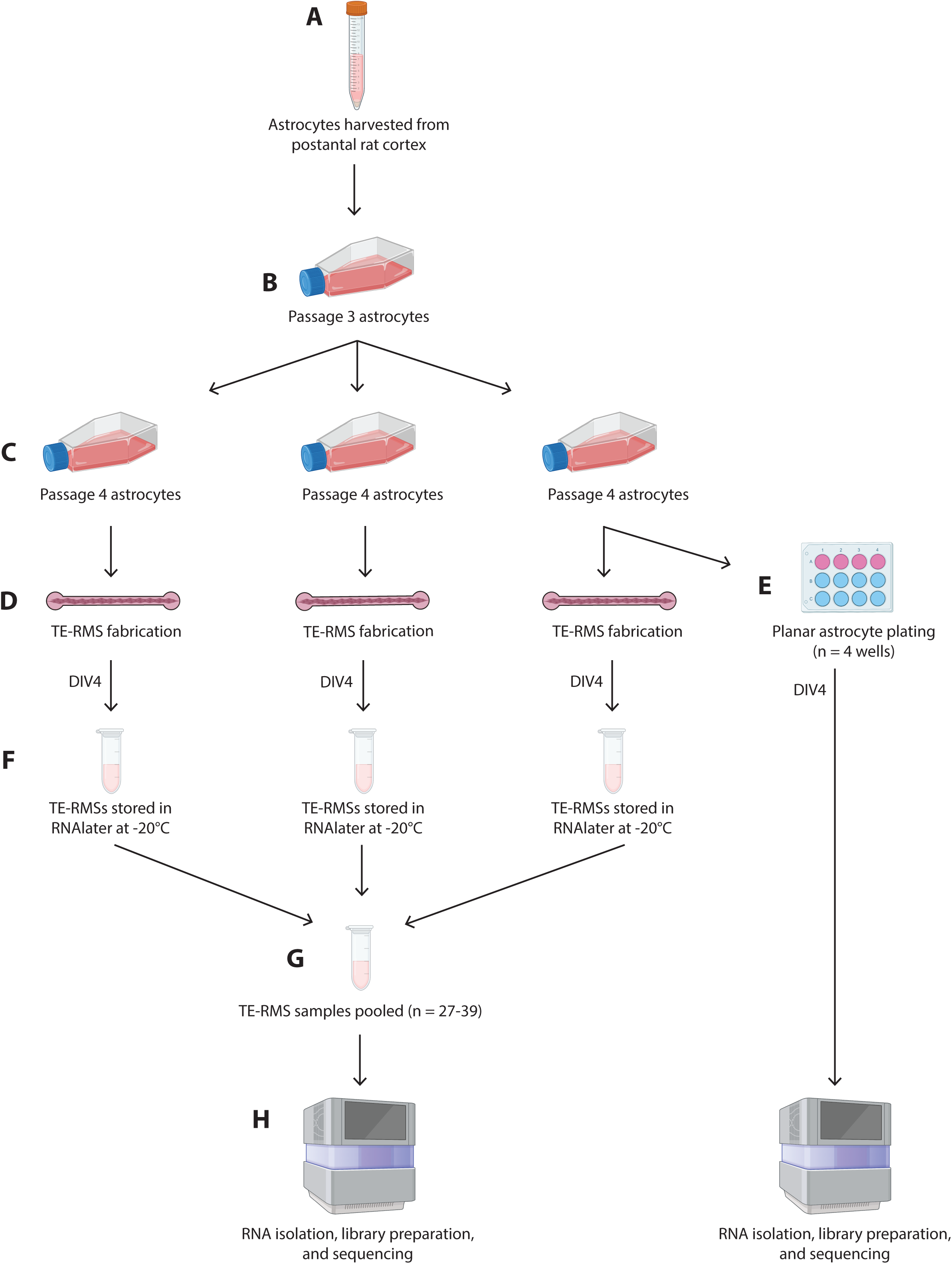
Experimental timeline. Astrocytes were harvested from the postnatal day 0-1 rat cortex (A). Cells were cultured in T75 flasks and purified through passaging. Passage 3 astrocytes (B) were split into 3 separate passage 4 sister flasks (C). 3 sequential TE-RMS fabrication days were required to generate a sufficient quantity of TE-RMS microtissues for RNA sequencing, with 1 T75 flask required for each fabrication day. On the third sequential day of TE-RMS fabrication, planar astrocytes were also plated in 4 wells of a 12-well plate (E). At DIV4 following TE-RMS fabrication, bundled microtissues were extracted from their microchannels, placed in RNAlater, and stored at −20°C (F). Preserved TE-RMS microtissues were pooled into one single sample (G). RNA was simultaneously extracted from DIV4 TE-RMS samples (frozen; n = 27-39 microtissues) and DIV4 planar astrocyte samples (fresh; n = 4 wells) (H). The entire process represented in this flow chart was repeated 3 times (defined as Run 1, Run 2, and Run 3), facilitating RNA sequencing on 3 distinct TE-RMS and 3 distinct planar astrocyte samples. Astrocytes were harvested from different animals for each repetition.

### Fabrication of molds and hydrogel microchannels

Fabrication of molds and hydrogel microchannels has been recently described in detail by our group (38). Briefly, microchannels were fabricated with custom stamps containing 9 channels arranged in a 3-by-3 grid with wells to connect media away from the channels (Figure 1A). Channels were 4.6 mm long, 0.3 mm wide, and 1.4 mm deep (Figure 1B). The desired channel design was reversed to create printable stamps that would create the channels when stamped into agarose. Stamps were printed in high-temperature resin at the University of Pennsylvania Libraries’ Holman Biotech Commons. Following printing, stamps were carefully removed from their supports, cleansed with de-ionized water, and autoclaved. Agarose (Sigma #A9539-500G) and phosphate buffered saline (Gibco #14190136) were mixed to create a 3% weight by volume agarose solution that was boiled until it was completely transparent and devoid of bubbles. Agarose was pipetted into 60 mm sterile dishes and spread evenly over the bottom of the dishes prior to stamps being placed into the hot agarose. Once the agarose had hardened, the stamps were removed. Three mL of PBS was added to each dish to keep the molds hydrated.

### Fabrication of tissue-engineered rostral migratory streams and planar astrocyte cultures

Fabrication of TE-RMSs in hydrogel microchannels has been recently described in detail by our group (38). Briefly, PBS was removed from microchannel molds with a glass Pasteur pipette and vacuum. Channels (Figure 1A) were seeded with cold rat tail type 1 collagen (Advanced BioMatrix #5153) diluted in neutralization buffer containing 50% TE-RMS culture media [a base of Neurobasal media supplemented with 2% B27 (Invitrogen #12587010), 0.25% L-glutamine (Gibco #35050061), 1% G5 supplement (Gibco #17503012), and 1% Penicillin-Streptomycin (Gibco #15140122)], 14.1% cell culture water (Corning #25055CM), 0.5X minimum essential media (Gibco #11430030), 25 mM HEPES (Gibco #15630080), 26.2 mM NaHCO3 (Gibco #25080094), and 0.5X G5 supplement (Gibco #17503012). Dishes were placed in an incubator at 37° C and 5% CO_2_ for around 2 hours until the collagen completely polymerized to coat the inner walls of the microchannels. During this collagen polymerization period, a flask of 80% confluent astrocytes was passaged with 0.25% trypsin-EDTA (Gibco #25200056) and cells were re-suspended in TE-RMS culture media at a density of 2.5 million cells/mL. Following complete collagen polymerization in the microchannels, astrocytes were seeded into the channels and dishes were returned to incubate at 37° C and 5% CO_2_. One hour later, dishes were flooded with TE-RMS culture media and returned to incubate at 37° C and 5% CO_2_. Under these conditions, astrocytes bundled together with collagen to self-assemble into TE-RMSs (Figure 1C-E). Three sequential TE-RMS fabrication days were required to generate a sufficient quantity of TE-RMS microtissues for RNA sequencing, with 1 T75 flask required for each fabrication day (Figure 2C-D). TE-RMSs at DIV4 after plating were used for all experiments (Figure 2D,F). For planar astrocyte samples, flasks of 80% confluent astrocytes were passaged with 0.25% trypsin-EDTA (Gibco #25200056). Cells were then resuspended in TE-RMS culture media and were plated on top of polymerized 1 mg/mL collagen in 12-well plates at a density of 2.5 million cells/mL in TE-RMS culture media. Planar cultures at DIV4 after plating were used for all experiments (Figure 2E).

### Immunocytochemistry

For fluorescent imaging, cultures were fixed with 4% paraformaldehyde for 30 minutes at room temperature. Planar astrocyte samples were fixed and stained directly on the coverslips on which the cells were grown. For TE-RMS samples, a glass coverslip was coated with 0.002% poly-l-lysine (Sigma #P4707), incubated for 2 hours at 37° C, rinsed 3 times with cell culture water (Corning #25055CM), and left to dry. A small quantity of freshly made collagen (see section detailing fabrication of tissue-engineered rostral migratory streams and planar astrocyte cultures) was placed on the coverslip and a TE-RMS was carefully extracted from its channel and laid onto the collagen-coated coverslip. Collagen was allowed to dry for 2-3 minutes prior to fixation. Following rinsing with PBS, cultures were permeabilized with 0.3% triton X-100 and blocked with 4% normal horse serum for one hour at room temperature. Following rinsing with PBS, TE-RMS and planar astrocyte cultures were incubated in goat anti-glial fibrillary acidic protein (GFAP) (1:1000) (Abcam #ab53554, RRID: AB_880202) overnight at 4° C. Cultures were then rinsed and incubated in donkey anti-goat 568 (1:500) (Thermo Fisher Scientific #A-11057, RRID: AB_2534104) and Hoechst solution (1:1000) (Thermo Fisher Scientific #H3570) in the dark for two hours at room temperature. Following secondary staining, cultures were rinsed with PBS, rinsed once with deionized water, then mounted onto glass slides with fluoromount G. The edges of the glass slides and coverslips were sealed with nail polish and allowed to dry in the dark prior to being stored at 4° C.

### Imaging

Phase images were obtained using a Nikon Inverted Eclipse Ti-S microscope with digital image acquisition using a QiClick camera interfaced with Nikon Elements Basic Research software (4.10.01). Fluorescent images were obtained using a Nikon A1Rsi Laser Scanning Confocal microscope with an x60 or x100 objective (CFI Plan Apo Lambda x60 Oil, n.a. 1.40; x100 Oil, n.a. 1.45).

### RNA extraction and RNA Sequencing

At DIV4 after fabrication, TE-RMSs were visualized with a phase microscope and only microtissues that were perfectly bundled were selected for RNA extraction. Bundled TE-RMSs were carefully extracted from their microchannels, placed immediately into RNAlater Solution (Invitrogen #AM7020), and stored at −20°C (Figure 2F). Planar astrocyte cultures were visualized with a phase microscope to ensure culture integrity prior to RNA extraction.

Preserved TE-RMS microtissues were pooled into one single sample (Figure 2G). Total RNA was extracted from TE-RMS samples and planar astrocyte samples simultaneously by the RNeasy Plus Universal Kit (73404, Qiagen, Hilden, Germany) (Figure 2H). The entire process was repeated 3 times (defined as Run 1, Run 2, and Run 3), facilitating RNA sequencing on 3 distinct TE-RMS and 3 distinct planar astrocyte samples. Quality of RNA was assessed via Bioanalyzer (Agilent, Santa Clara, CA) automated electrophoresis, ensuring an RNA Integrity Number above 8.0 and the presence of 18S and 28S rRNA bands. Extracted RNA was reverse transcribed to complementary DNA and the sequencing library prepared using the SMART-Seq mRNA LP (634768, Takara Bio Inc., Kusatsu, Shiga, Japan). Sequencing was performed on an Illumina NextSeq 2000 (San Diego, CA) to generate 100 base pair single reads and approximately 20M reads per sample (Figure 2H).

### Data Analysis

Fastp was used to qc and trim the sequencing fastq files and Salmon was used to count the trimmed data against the transcriptome defined in Ensembl v111, which was built on the genome mRatBN7.2 (42,43). Several Bioconductor (v3.18) packages in R (v4.33) were used for annotation and analysis (44). The transcriptome count data was annotated and summarized to the gene level with tximeta and biomaRt (45,46). Count data was analyzed in a paired structure so that TE-RMS microtissues and planar astrocytes from the same run were considered paired. Paired count data was analyzed with Principle Component Analysis (PCA) analysis and plots were generated with PCAtools (47). Heatmaps were done with ComplexHeatmap (48).

Normalizations and statistical analyses were done with DESeq2 (49). False discovery rate was calculated with the Benjamini-Hochberg procedure. Differentially expressed genes were defined as those with an adjusted p-value < 0.05. GSEA pathway analysis was done using the Molecular Signatures Database (MSigDB) on gene sets related to cytoskeleton and nuclear structure (50,51). Analysis was performed on annotated genes only. Significance threshold was set to p < 0.05.

## Results

To fabricate TE-RMSs, grids of hydrogel rectangular microchannels (Figure 1A-B) were dried and coated with extracellular matrix. Following complete polymerization and drying of extracellular matrix, the microchannels were seeded with a dense suspension of astrocytes that bundled to form longitudinally-aligned TE-RMSs (Figure 1C). This fabrication process during which astrocytes self-assemble into a three-dimensional TE-RMS induced profound morphological changes in the astrocytes compared to counterpart astrocytes in planar culture. Specifically, cytoskeletal rearrangement and nuclear elongation were clearly evident in TE-RMS astrocytes. Phase microscopy and high magnification confocal imaging revealed elongated cell nuclei and bidirectional intermediate filament processes in TE-RMSs (Figure 1E) while planar astrocyte samples had round nuclei and intermediate filament processes spread in all directions (Figure 1F-H).

To begin evaluating transcriptomic changes in TE-RMS astrocytes compared to planar astrocytes, we pooled samples for each run, extracted total RNA, assessed RNA quality, then conducted RNAseq. From the RNAseq output, we first generated a heatmap with hierarchical clustering using normalized count data of a subset of genes (Figure 3). This heatmap served as a high level overview of highly expressing genes and the hierarchical clustering tree diagram helped visualize the similarities between specimens. The rows of genes in the upper half of the heatmap had the highest normalized counts for TE-RMS astrocytes while the genes in the lower half had the highest counts for planar astrocytes. Additionally, the TE-RMS sample columns had many rows of highly expressing genes for all 3 runs, potentially indicating genes for future analysis. Through hierarchical clustering, we noted the clustering of all 3 TE-RMS runs with each other and the clustering of all 3 planar astrocyte runs with each other. This suggested that the gene expression patterns of TE-RMSs were more similar to each other compared to planar astrocytes.

**Figure 3.**
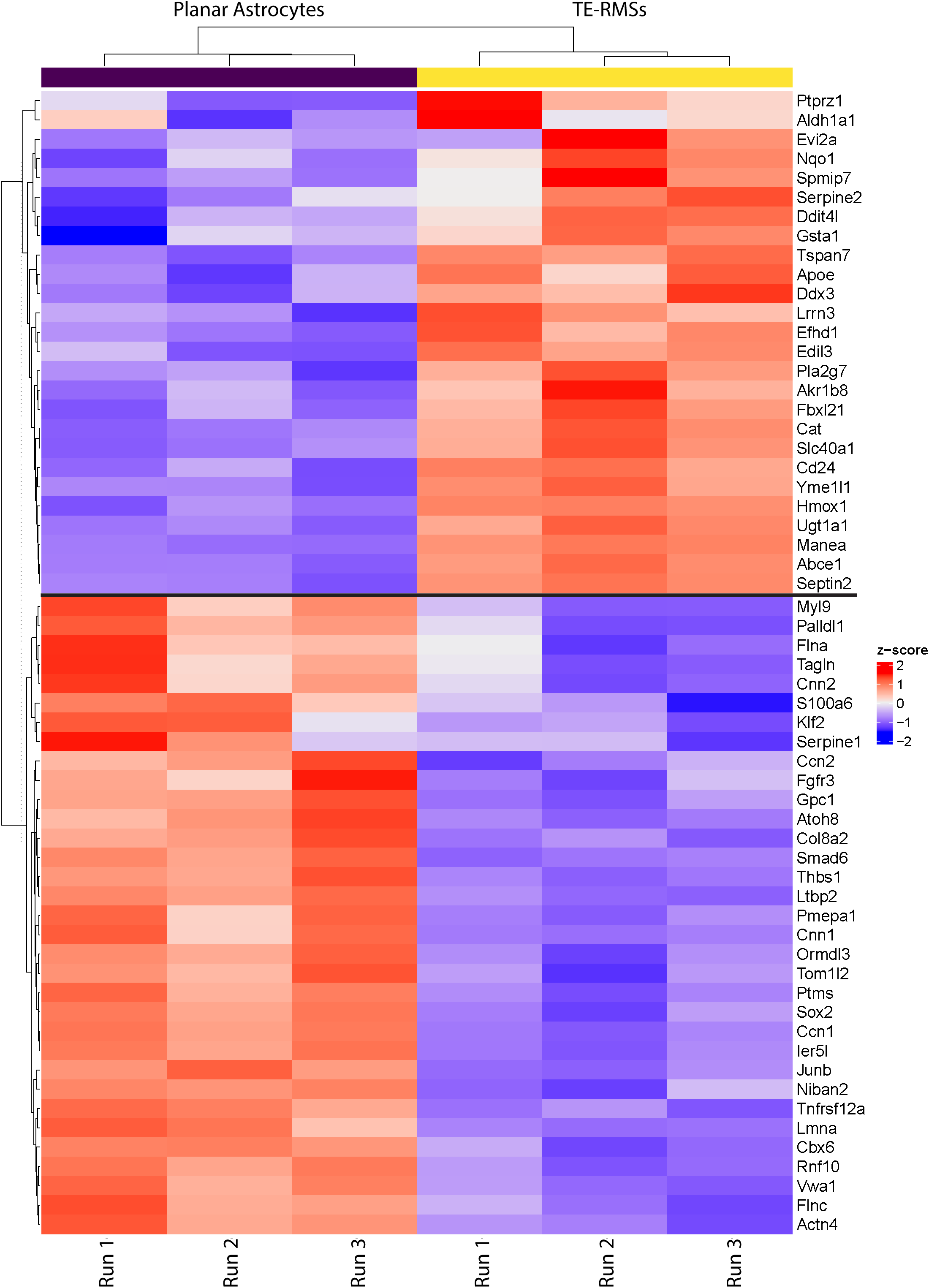
Heatmap with hierarchical clustering. Using normalized count data, we plotted a subset of genes in the upper portion of the graph that had the highest normalized counts for TE-RMSs, while a subset of genes in the lower portion of the graph had the highest normalized counts in planar astrocytes. Specimens are represented by columns and each gene is represented by a row. Red indicates high expression and blue indicates low expression. Through the tree diagram, hierarchical clustering demonstrated that TE-RMSs closely associate with each other and planar astrocytes closely associate with each other. Additionally, there are many rows of highly expressed genes in the TE-RMS columns that may indicate gene candidates for future studies.

We continued our exploratory data analysis through PCA, which processed our large transcriptomic data set and transformed it into smaller sets that still contain the majority of the information. The result of this reduced data set complexity was a simplistic overview that demonstrated clear separation between our experimental groups. The first two PCs of the gene expression data explained over 70% of the variance (Figure 4). PC1 explained 51.49% and PC2 explained 19.76%. Along PC1, we saw a clear separation in our experimental groups as TE-RMS samples clustered on the right of the axis, while planar astrocytes clustered on the left side of the axis. Interestingly, PC2 separated our samples according to experimental run as TE-RMS and planar astrocyte specimens from Run 1 clustered with each other, as Run 2 specimens clustered with each other, and Run 3 specimens clustered with each other. This indicated some gene expression differences between sample runs; however, this PC2 percent variation was smaller and less influential than PC1. Overall, our PCA results suggested the transcriptomic expression data consolidated into PCs can differentiate TE-RMS astrocytes from planar astrocytes.

**Figure 4.**
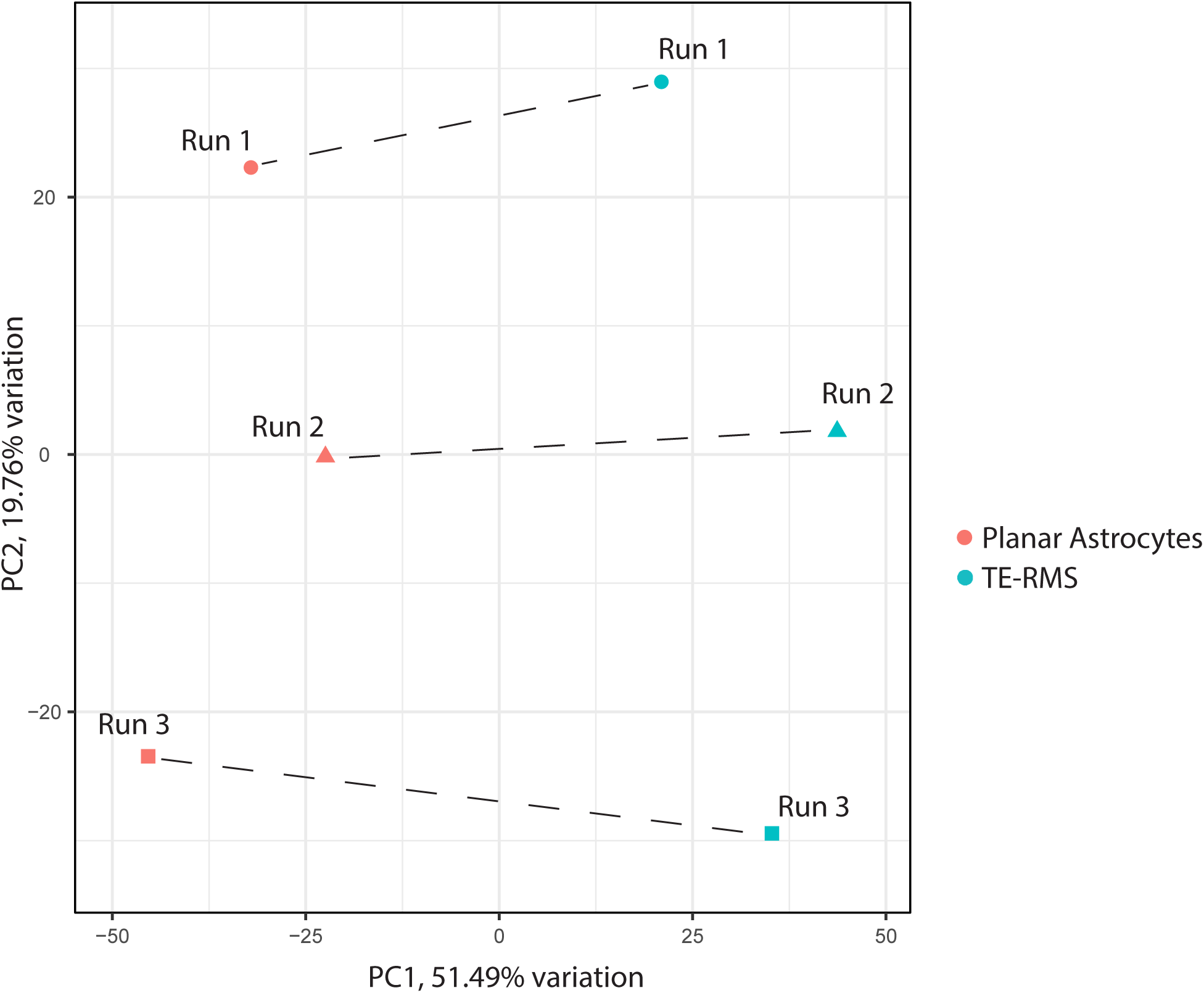
Principal component analysis. The first 2 principal components (PC) are labeled followed by the percentage of variance explained by each PC. We visualized the close clustering of TE-RMS astrocytes versus planar astrocytes along PC1, and the clustering of experimental runs along PC2. These PC analysis results suggested that PCs can differentiate TE-RMS astrocytes from planar astrocytes, as well as experimental run to a lesser extent.

We performed differential expression (DE) analysis to determine which gene expression differences were statistically significant between TE-RMS astrocytes and planar astrocytes. Compared to planar astrocyte expression, 4008 genes were differentially expressed in TE-RMS astrocytes, with 2076 downregulated genes and 1932 upregulated genes (Figure 5). To examine only the most impactful genes, we applied a cutoff of ±3 fold change (log2), with 256 genes downregulated and 91 genes upregulated. Tables below present the top ten downregulated (Table 1) and upregulated (Table 2) genes. Additionally, we examined our DE analysis results for *robo2* and *ezrin* expression as we have previously reported that the endogenous RMS and TE-RMS astrocytes have enriched Robo2 and Ezrin proteins compared to non-RMS astrocytes *in vivo* or planar astrocyte cultures *in vitro* (12). DE analysis revealed a non-significant increase (1.17 log2 fold) in Robo2 expression (p = 0.0791) and a significant increase (0.79 log2 fold) in Ezrin expression (p < 0.0001).

**Figure 5.**
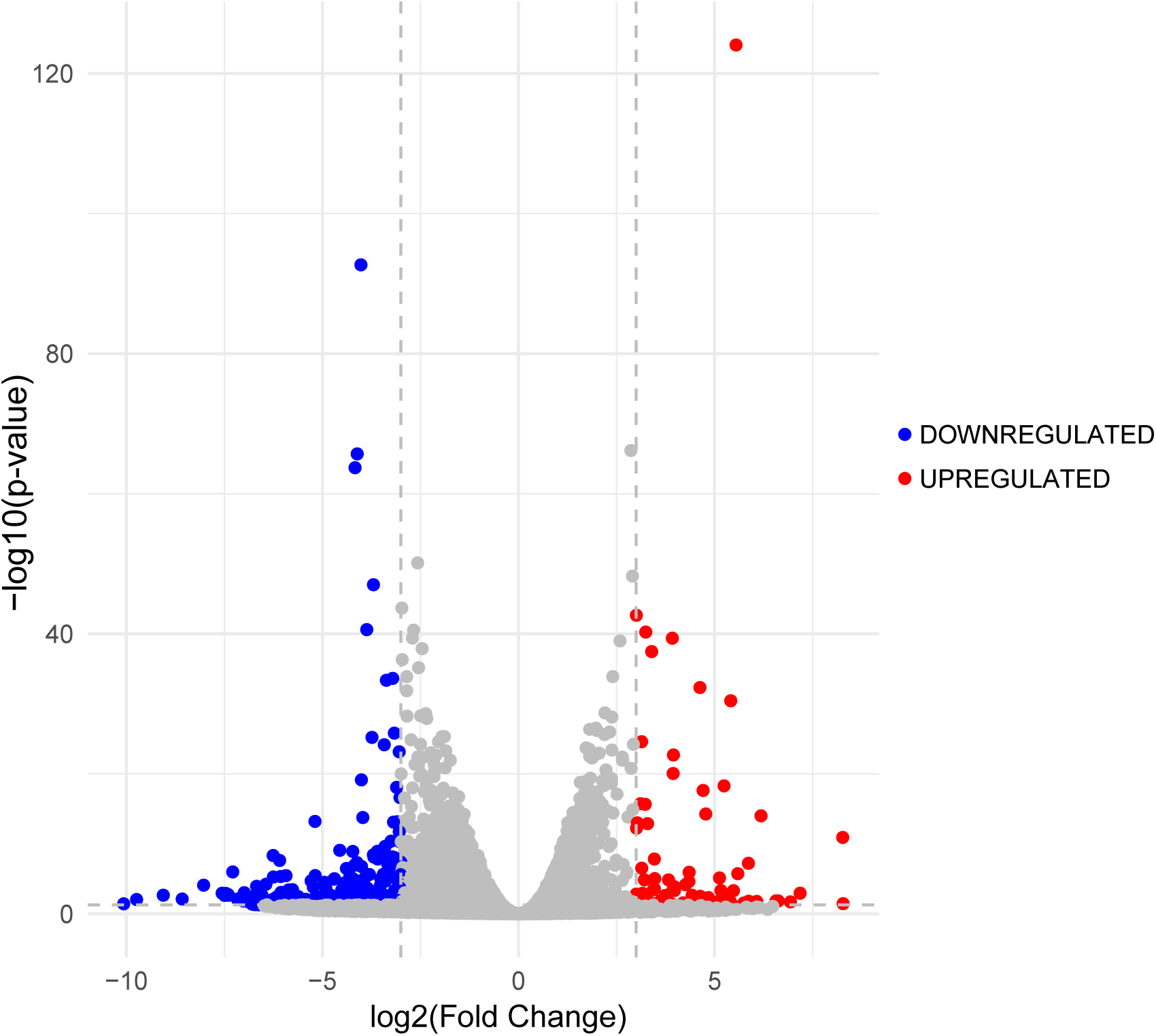
Differentially expressed genes in TE-RMS compared to planar astrocytes. The volcano plot depicted each gene’s -log_10_(p-value) and log_2_ fold change. To examine only significant and highly impactful genes, the horizontal dashed line indicated false discovery rate adjusted p-value < 0.05 and the vertical dashed lines indicated ±3 fold change. 256 downregulated genes and 91 upregulated genes met these criteria.

**Table 1.**
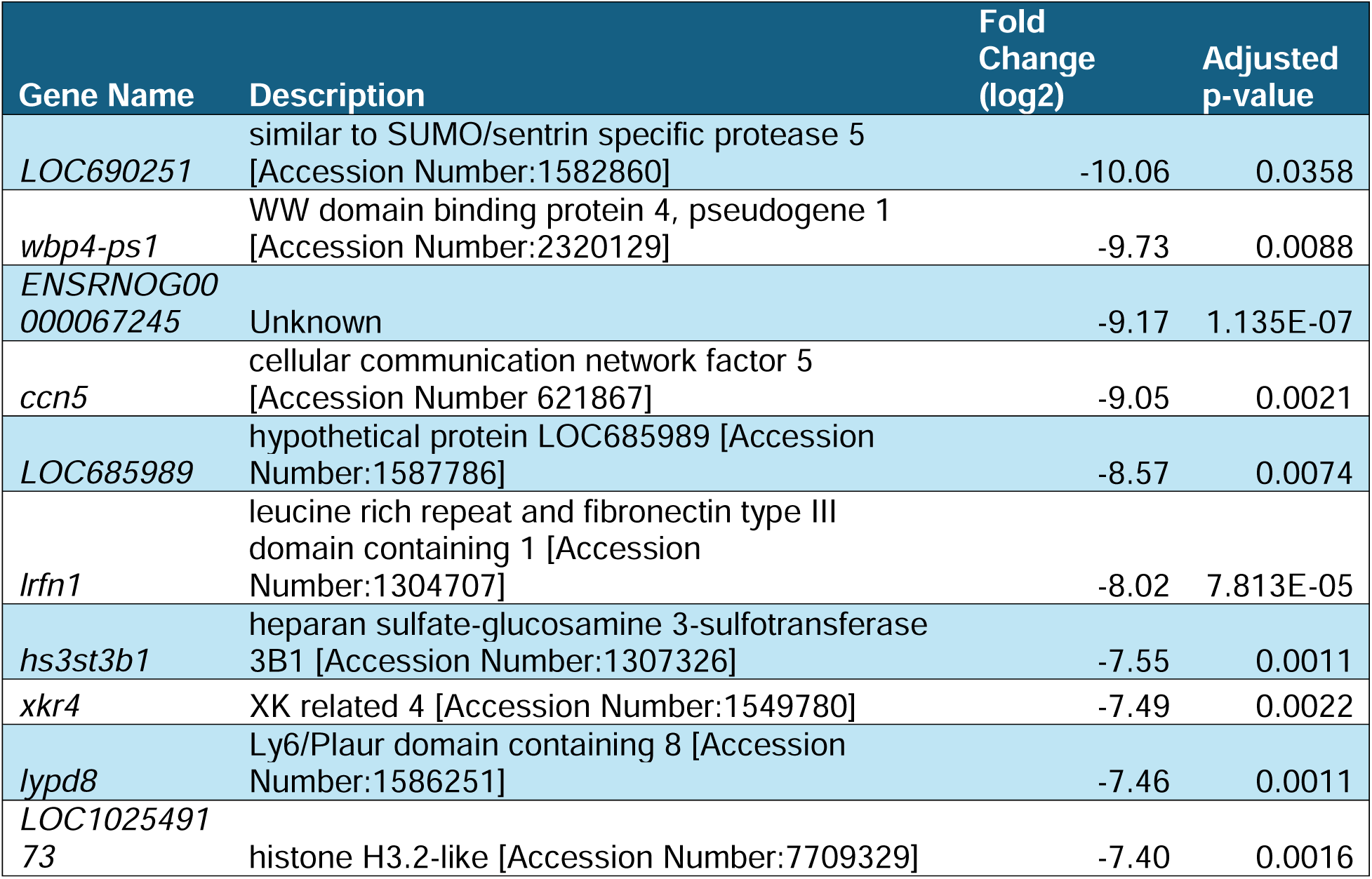
Top ten downregulated genes with the greatest fold change (log2).

**Table 2.**
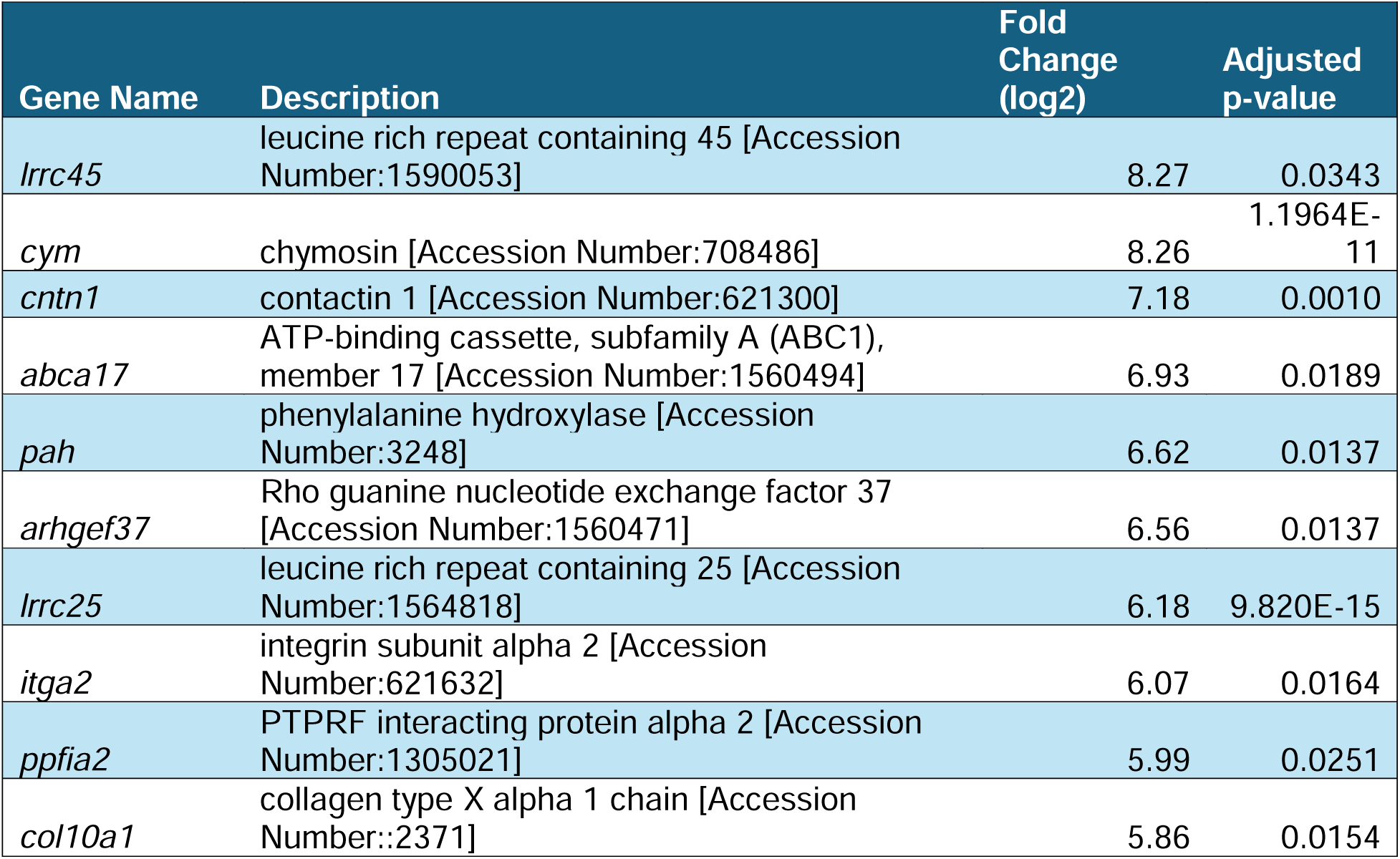
Top ten upregulated genes with the greatest fold change (log2).

Our previous research has shown that the unique morphology of TE-RMS astrocytes mimicked the unique cytoskeletal and nuclear architecture of the endogenous rat RMS (37). Therefore, we conducted gene set analysis on cellular component sets related to cytoskeleton (Figure 6) and nuclear (Figure 7) structure to identify gene expression changes that may underlie these morphological changes. Using the molecular signatures database, we assessed annotated gene sets for the following: Cytoskeleton, Actin Cytoskeleton, Actin Filament, Microtubule Cytoskeleton, Polymeric Cytoskeleton, Septin Cytoskeleton, Spectrin Associated Cytoskeleton, Nuclear Body, Nuclear Chromosome, Nuclear Envelope Lumen, Nuclear Inner Membrane, Nuclear Lamina, Nuclear Membrane, Nuclear Outer Membrane, Nuclear Periphery, and Nuclear Pore.

**Figure 6.**
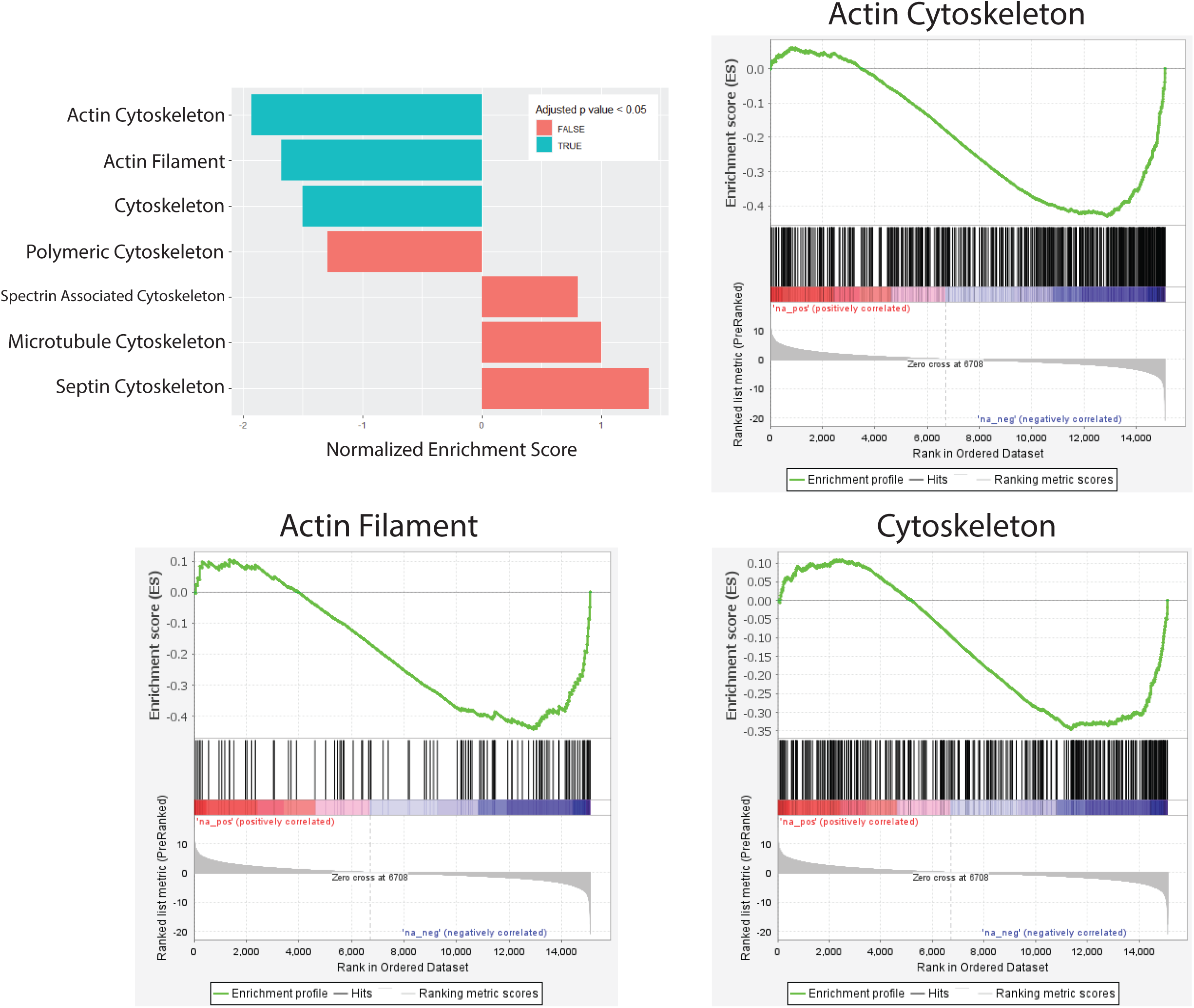
Cytoskeleton gene set analysis. Gene set analysis was conducted on cellular components related to cytoskeleton structure using the molecular signatures database annotated gene sets. Cellular components were significantly enriched for Actin Cytoskeleton (454 genes, NES = −1.930, adjusted p-value < 0.0001), Actin Filament (106 genes, NES = −1.681, adjusted p-value = 0.0034), and Cytoskeleton (332 genes, NES = −1.500, adjusted p-value = 0.0142). Graphical representations of the rank metric score and the running enrichment score are presented for Actin Cytoskeleton, Actin Filament, and Cytoskeleton gene sets.

**Figure 7.**
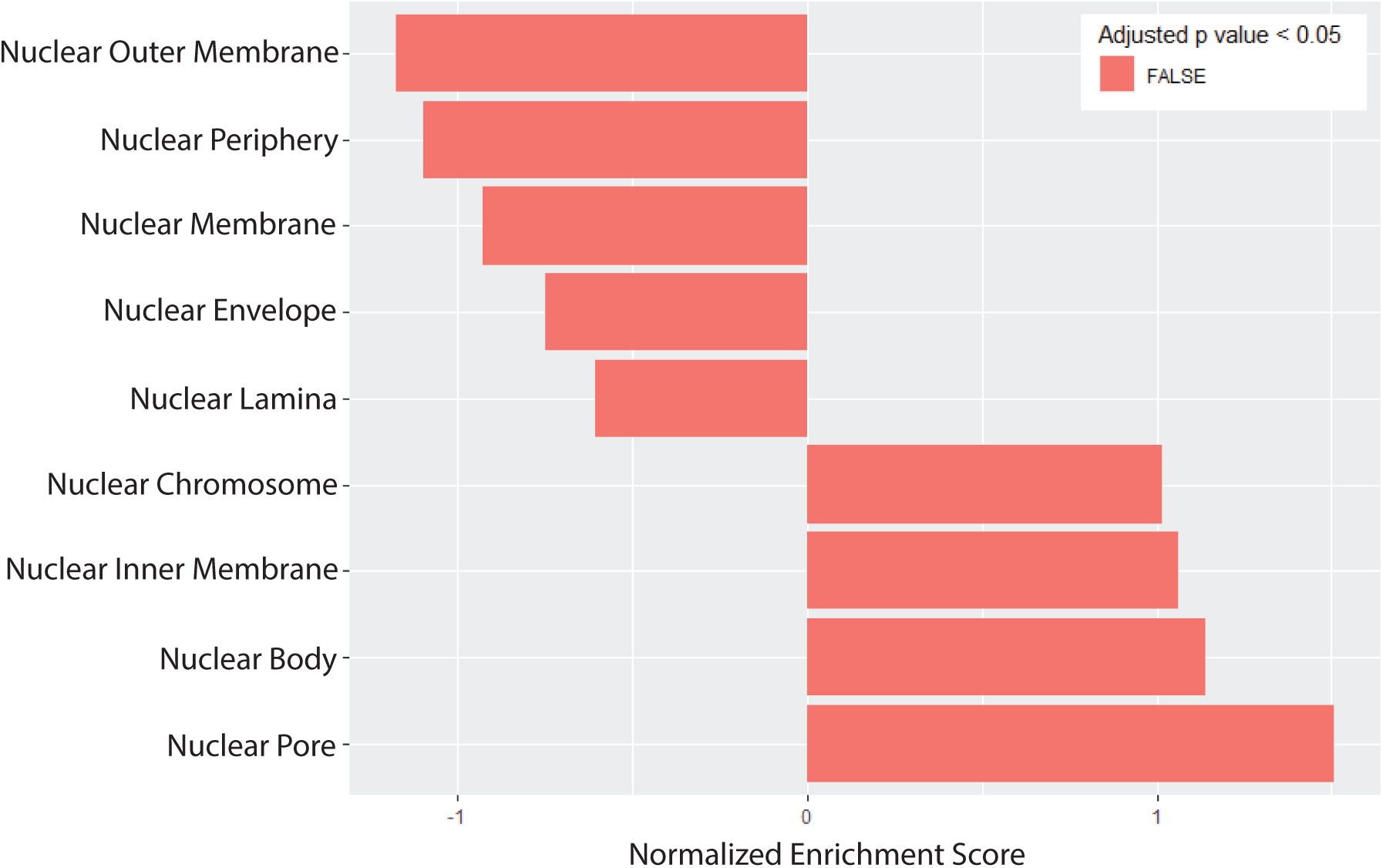
Nuclear gene set analysis. Gene set analysis was conducted on cellular components related to nuclear structure using the molecular signatures database annotated gene sets. No gene set was significantly enriched.

From the p-values of our DE analysis, we ranked all our genes and generated a local statistic, called a rank metric score. This rank metric allowed us to sort our genes so that upregulated genes with small p-values were at the top of the list and downregulated genes with small p-values were at the bottom of the list. We also generated a running enrichment score for each gene by starting at the top of our ranked gene list and adding to the running sum if the gene is a member of the gene set of interest or subtracting if the gene is not in that gene set. Finally, the enrichment score was the largest value (positive or negative) achieved during the running sum. This score is normalized depending on the gene set size and labeled as our normalized enrichment score (NES), which allowed direct comparison of all gene sets analyzed. We calculated cellular components that were significantly enriched for Cytoskeleton (332 genes, NES = −1.500, adjusted p-value = 0.0142) (Table 3), Actin Cytoskeleton (454 genes, NES = −1.930, adjusted p-value < 0.0001) (Table 4), and Actin Filament (106 genes, NES = −1.681, adjusted p-value = 0.0034) (Table 5) (Figure 6). No gene set was significantly enriched for nuclear structure (Figure 7).

**Table 3.**
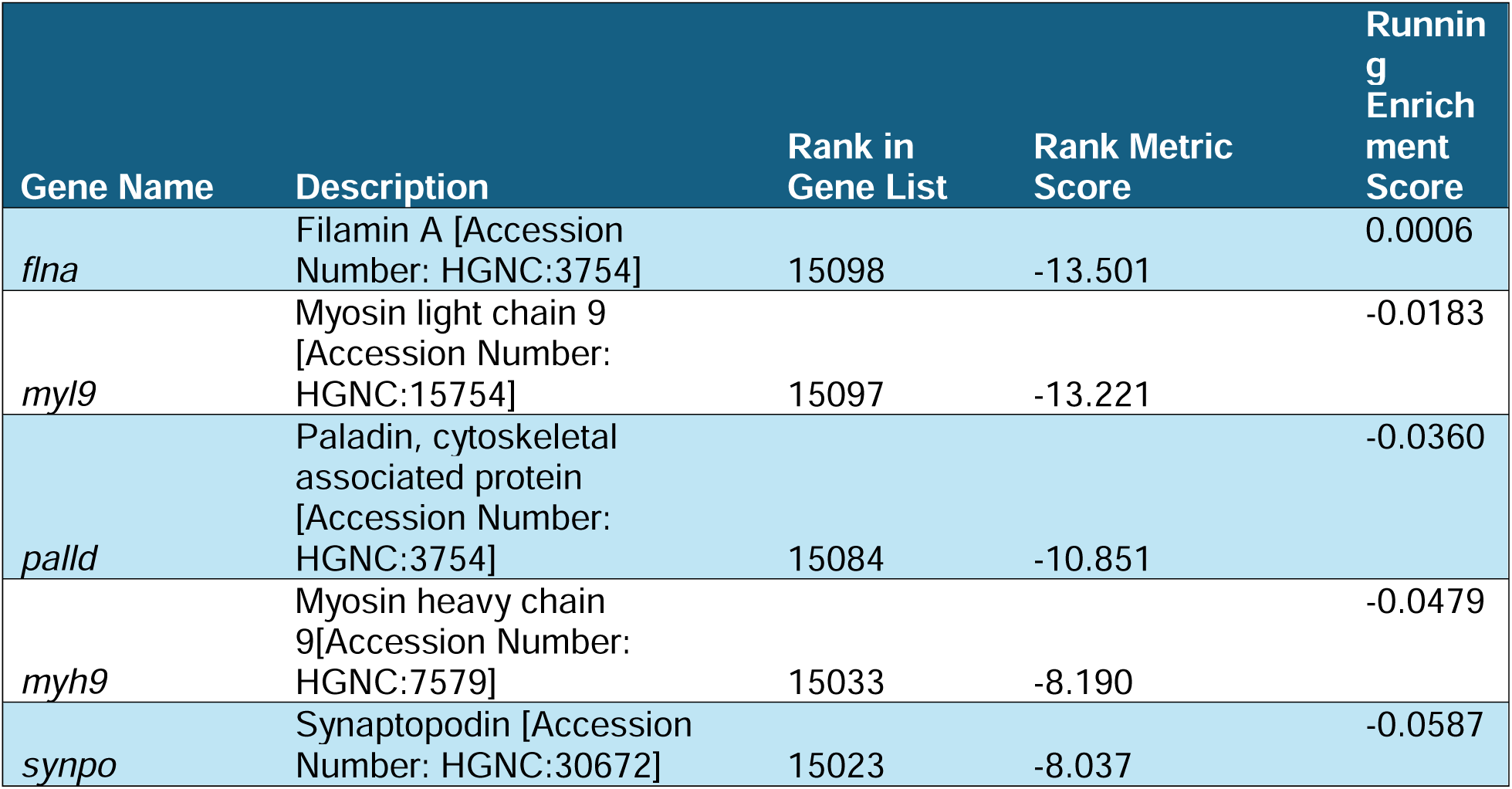
Top 5 over-represented genes in the Cytoskeletal gene set that are downregulated.

**Table 4.** Top 5 over-represented genes in the Actin Cytoskeletal gene set that are downregulated.

| Gene Name | Description | Rank in Gene List | Rank Metric Score | Running Enrichment Score |
| --- | --- | --- | --- | --- |
| *cnn1* | Calponin 1 [Accession Number: HGNC:2155] | 15101 | -13.986 | 0.0004 |
| *flna* | Filamin A [Accession Number: HGNC:3754] | 15098 | -13.501 | -0.0132 |
| *myl9* | Myosin light chain 9 [Accession Number: HGNC:15754] | 15097 | -13.221 | -0.0265 |
| *actn4* | Actinin Alpha 4 [Accession Number: HGNC:166] | 15085 | -10.932 | -0.0388 |
| *palld* | Paladin, cytoskeletal associated protein [Accession Number: HGNC:3754] | 15084 | -10.851 | -0.0495 |

**Table 5.** Top 5 over-represented genes in the Actin Filament gene set that are downregulated.

| Gene Name | Description | Rank in Gene List | Rank Metric Score | Running Enrichment Score |
| --- | --- | --- | --- | --- |
| *flna* | Filamin A [Accession Number: HGNC:3754] | 15098 | -13.501 | 0.0006 |
| *palld* | Paladin, cytoskeletal associated protein [Accession Number: HGNC:3754] | 15084 | -10.851 | -0.0480 |
| *actn1* | Actinin Alpha 1 [Accession Number: HGNC:163] | 15042 | -8.538 | -0.0851 |
| *lmod1* | Leiomodin 1 [Accession Number: HGNC:6647] | 15019 | -7.929 | -0.1149 |
| *tpm2* | Tropomyosin 2 [Accession Number: HGNC:12011] | 15000 | -7.515 | -0.1428 |

## Discussion

The TE-RMS is a tissue engineered biological scaffold comprising aligned astrocytes designed to facilitate NPC migration. Our previous work described morphological changes to astrocytes during their self-assembly into the TE-RMS, specifically cytoskeletal remodeling and nuclear elongation (37). In the current study, we sought to examine the genetic underpinnings of these morphological changes through RNA sequencing. We found 256 downregulated genes and 91 upregulated genes with over a 3-fold change in TE-RMS astrocytes compared to planar astrocytes. Additionally, we hypothesized that TE-RMS astrocytes would have significant changes to cytoskeletal and nuclear gene sets compared to planar astrocytes owing to their altered cytoarchitectural and nuclear morphology. Using gene set enrichment analysis, we found significant enrichment for cytoskeleton, actin cytoskeleton, and actin filament gene sets. However, we did not find significant enrichment for any nuclear structure-related gene sets, so we are compelled to partially reject our hypothesis.

RNA sequencing allowed us to map the complex cellular responses and characterize the molecular phenotype of the unique TE-RMS astrocytes. Our higher level analyses – heatmaps with hierarchical clustering and PCA – demonstrated that TE-RMS astrocytes had observationally distinct expression profiles compared to planar astrocytes. PCA results appeared to partially differentiate between TE-RMS astrocytes and planar astrocytes and, to a lesser degree, the experimental run from which the astrocytes were isolated.

Yet it is through the differential expression analysis that we determined the individual genes that were significantly different between our experimental groups. One of our significantly downregulated genes was LOC690251 (−10.06 fold). LOC690251 is similar to a SUMO/sentrin specific protease 5, which breaks down the SUMO proteins involved in many post-translational modifications and may be required for cell division (52,53). Downregulation of LOC690251 suggests that astrocyte cell division within the TE-RMS may be minimized, possibly due to the astrocytes prioritizing alignment, cell-cell adhesions, and expression of functional proteins. *ccn5* (−9.05 fold) is a cellular communication network factor relevant to a connective tissue growth factor family and involved in the regulation of cancer progression (54). *lrfn1* (−8.02 fold), a family of adhesion molecules also known as synaptic adhesion-like molecules (SALM), is involved in neurite outgrowth and synapse formation in neurons, however its role in astrocytes has not been established (55). Downregulation of *ccn5* and *lrfn1* may alter astrocyte adhesion to each other and the ECM, which may benefit the TE-RMS as astrocytes need to align in a bidirectional, tightly-coupled pathway with specific adhesion patterns as opposed to the omnidirectional adhesion and communication typical of stellate astrocytes within the brain parenchyma.

One of the significantly upregulated genes included *lrrc45* (8.27 fold), which connects centrosomes during interphase of the cell cycle and organizes microtubules (56). Centrosome cohesion may affect numerous cellular processes including cell polarity, motility, and transport - suggesting an important influence on TE-RMS astrocyte motility, bipolar shape, and ability to divide if needed (57). Another significantly upregulated gene is *cntn1* (7.18 fold), a cell surface adhesion molecule gene. When overexpressed in GFAP-expressing glioblastomas, the molecule creates a repellent effect on glioma cells, which may also play a role in TE-RMS cell organization and alignment (58). *cntn1* is also implicated in inflammation disorders and may facilitate crosstalk between astrocytes and microglia (59). *pah* (6.62 fold) encodes for an enzyme that processes phenylalanine and alters astrocyte and microglia morphology in a *pah* variant-spliced mouse model (60). Future experiments should assess if *pah*, along with *lrrc45* and *cntn1*, contribute to the bidirectional process filaments characteristic of TE-RMS astrocytes.

We also examined our DE analysis results for *robo2* and *ezrin* expression. Astrocytic Robo2 receptors ensure proper neuroblast migration in the RMS through chemorepulsion, while ezrin is a membrane-cytoskeletal protein expressed at high levels in RMS astrocytes (23,61,62). We also previously reported that the astrocytes in the RMS *in vivo* and TE-RMS astrocytes *in vitro* both have more Robo2 and Ezrin than non-RMS astrocytes *in vivo* or planar astrocyte cultures *in vitro* (12). Interfering with the function of these proteins impedes neuroblast migration through the RMS. In our current analyses, *ezrin* expression was significantly upregulated (p<0.0001), while *robo2* expression was upregulated but narrowly missed the threshold for statistical significance (p=0.0791). *ezrin* upregulation further validates our TE-RMS model with key characteristics of the endogenous rat RMS. While *robo2* expression was not statistically significant, other transcripts related to cell migration, such as *lrrc45* and *cntn1*, are greatly upregulated suggesting that TE-RMS astrocytes express factors believed to facilitate neuroblast migration. Additionally, Robo2 receptors may undergo post-translation modification to function optimally in the RMS, which would not be reflected in the current study’s approach. Future studies should examine Ezrin and Robo2 gene and protein-level function over time in TE-RMS astrocytes.

Unlike our DE analysis, our gene set enrichment analysis allowed us to examine expression changes in functionally-related gene sets. Using the molecular signatures database, we looked at all cellular component gene sets related to cytoskeleton and nuclear structure and found significantly enriched gene sets for cytoskeleton, actin cytoskeleton, and actin filament gene sets, while no nuclear structure gene sets were significantly enriched. This may suggest that while the angle of curvature in the agarose channels provided mechanical cues that signal gene expression changes related to cytoskeletal rearrangement, the nuclear elongation observed in RMS and TE-RMS astrocytes may result from cytoskeletal forces and not expression changes with nuclear structure genes. Additional studies will be needed to differentiate between expression changes contributing to cytoskeletal rearrangement and those that resulted from cytoskeletal elongation generating forces that act on the nucleus and potentially rearranging chromatin.

Following the enrichment analysis, which determined whether an *a priori* defined set of genes showed statistical significance between two phenotypes, we identified over-represented genes in each significant gene set (Table 6). These are genes that are present more than would be expected in our selected gene set data. The over-represented genes in the cytoskeletal gene set were downregulated and include *flna*, which encodes for an actin-binding protein filamin A and serves as a versatile molecular scaffold (63). *myl9* regulates myosin light chain, is highly expressed in astrocytes, and dramatically decreases after small molecule reprogramming of astrocytes into neurons (64). *palld* is involved in actin cytoskeleton organization and its expression levels vary in astrocytes depending on their morphology (65). *myh9* is also involved in actin-binding and crucial for adhesion and cell migration (66). Finally, *synpo* encodes synaptopodin, an actin-binding protein that is typically found in postsynaptic densities and dendritic spines in neurons, and may protect F-actin from disruption; however its role in astrocytes is unknown (67).

**Table 6.**
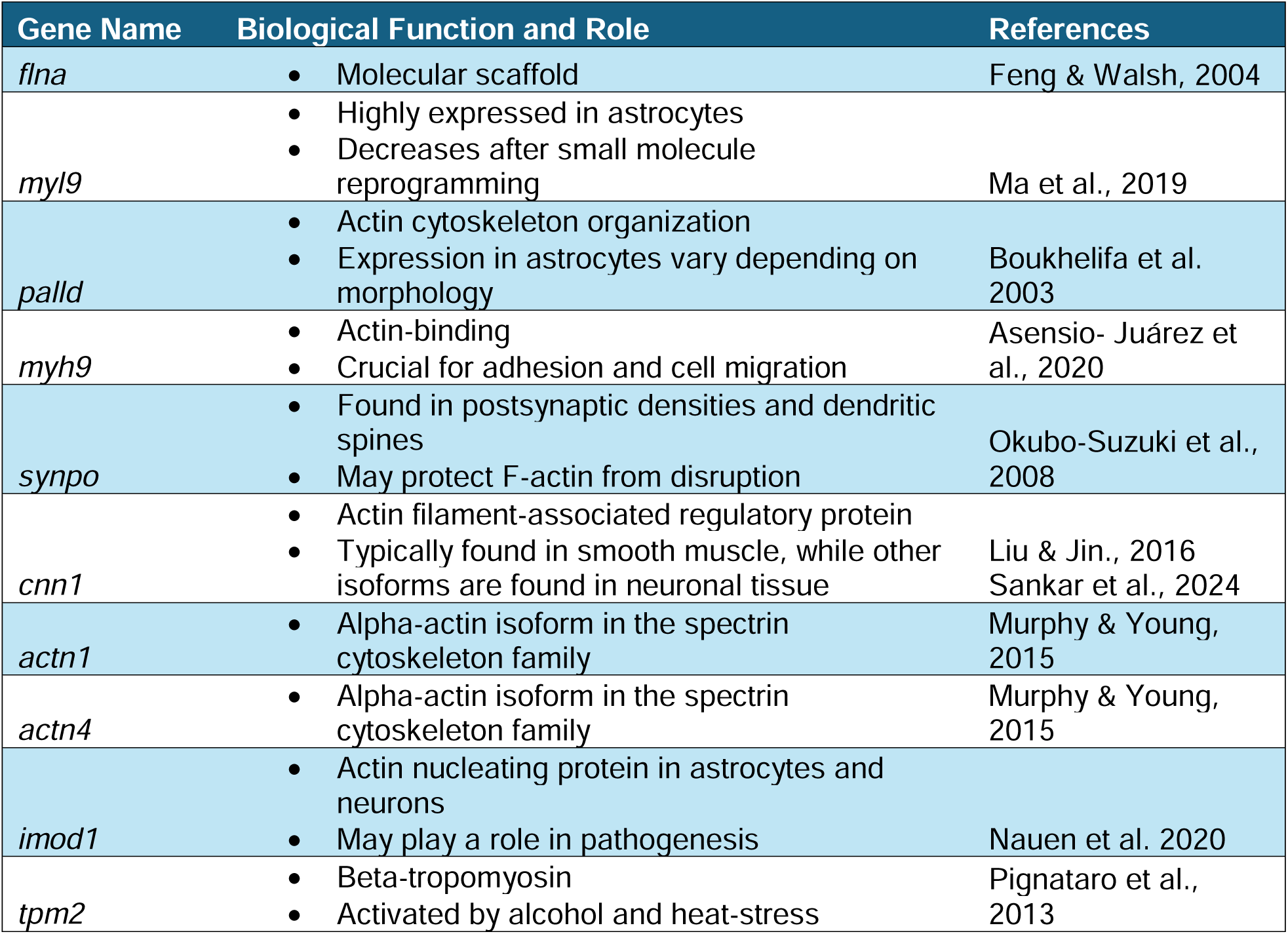
Function and role of over-represented genes in cytoskeletal and nuclear gene sets.

Within the actin cytoskeleton and actin filament gene sets, we also see *flna*, *myl9*, *palld* as over-represented genes that are downregulated. Additionally, *cnn1* encodes for calponin 1, an actin filament-associated regulatory protein. Calponin isoforms *cnn2* and *cnn3* are expressed in neuronal tissue, while *cnn1* is typically found in smooth muscle cells, so its role in the current study is unclear (68,69). *actn1* and *actn4* are alpha-actinin isoforms and belong to the spectrin gene superfamily, which is a group of cytoskeletal proteins (70). *lmod1* is an actin nucleating protein present in astrocytes and neurons, and may play a role in pathogenesis (71). *tpm2* encodes beta-tropomyosin and is activated by alcohol and heat-stress in astrocyte cultures (72).

Overall, these gene set analyses show the high degree of influence of structural protein-related genes that are downregulated in TE-RMS astrocytes. This is unsurprising as we have witnessed stark structural changes in TE-RMS astrocytes; planar astrocytes extend in all directions, while the aligned astrocytes of the TE-RMS take on a bidirectional morphology. Other literature has noted astrocyte cytoskeleton changes in aging astrocytes and during distress, conditions which produce atrophied or hypertrophied astrocytes (73). These morphologies are distinct from the bidirectional astrocytes of the TE-RMS. Furthermore, astrocytic studies specifically focused on impaired actin dynamics note reduced morphological complexity as well as decreased astrocyte infiltration, which results in diminished axonal pruning (74,75).

While the current study was primarily descriptive, the data generated are useful to generate testable hypotheses for future studies. For instance, future studies could genetically alter these actin-related genes to study the effects on cell morphology and function—particularly nucleo-cytoskeletal interactions. Future studies could also test the efficacy of manipulating these genes to enhance the fabrication or function of the TE-RMS. Cell migration assays using genetically altered TE-RMSs would inform us of the optimal TE-RMS cytoarchitecture to improve its ability to facilitate migration. Integrating proteomic analyses would allow us to verify the protein abundance of our DE genes and examine post-translational modifications.

Overall, the current study performed a detailed transcriptomic evaluation of a unique astrocyte-based engineered microtissue compared to planar counterparts. TE-RMS astrocytes demonstrated extensive actin-related gene expression changes paired with histologically observed cytoskeletal rearrangement and significantly altered nuclear shape. This study highlights the TE-RMS as a novel platform to investigate factors underlying cytoskeletal arrangement, nucleo-cytoskeletal interactions, astrocyte function, and cell migration. Our findings offer an array of genetic targets that can be manipulated to systematically test the underlying mechanisms and/or augment these functions to enhance the efficacy of these living regenerative scaffolds.

## Acknowledgements

The authors thank the University of Pennsylvania Libraries’ Holman Biotech Commons for 3D printed molds and the Penn Genomics and Sequencing Core (RRID:SCR_022382).

## Ethical Approval

All procedures were approved by the Institutional Animal Care and Use Committees at the University of Pennsylvania and the Michael J. Crescenz Veterans Affairs Medical Center and adhered to the guidelines set forth in the NIH Public Health Service Policy on Humane Care and Use of Laboratory Animals (2015).

## Authorship

Conceptualization, M.R.G., E.M.P., J.C.O., D.K.C.; methodology, M.R.G., E.M.P., A.D.G.E., E.N.K.; software, M.R.G.; validation, M.R.G.; formal analysis, M.R.G.; investigation, M.R.G.; resources, J.C.O., D.K.C.; data curation, M.R.G.; writing—original draft preparation, M.R.G.; writing—review and editing, M.R.G., E.M.P., J.C.O., D.K.C.; visualization, M.R.G., E.M.P.; supervision, J.C.O., D.K.C.; project administration, D.K.C.; funding acquisition, J.C.O., D.K.C. All authors have read and agreed to the published version of the manuscript.

